# Monocyte and Macrophage Subtypes as Paired Cell Biomarkers for Coronary Artery Disease

**DOI:** 10.1101/323287

**Authors:** Kathryn A. Arnold, John E. Blair, Jonathan D. Paul, Atman P. Shah, Sandeep Nathan, Francis J. Alenghat

## Abstract

**Background:** Monocytes and macrophages are central to atherosclerosis, but how they mark progression of human coronary artery disease (CAD) is unclear. We tested whether patients’ monocyte subtypes paired with their derived macrophage profiles correlate with extent of CAD.

**Methods**: Peripheral blood was collected from 30 patients undergoing cardiac catheterization, and patients were categorized as having no significant CAD, single vessel disease, or multivessel disease according to the number of affected coronary arteries. Mononuclear cells were measured for monocyte markers CD14 and CD16 by flow cytometry, and separate monocytes were cultured into macrophages over 7 days and measured for polarization markers CD86 and CD206.

**Results:** At baseline, patients with greater CAD burden were older with higher rates of statin use, whereas all other characteristics were similar across the spectrum of coronary disease. Non-classical (CD14^lo^CD16^hi^) and all CD16^+^ monocytes were elevated in patients with single vessel and multivessel disease compared to those without significant CAD (8.6% and 10.5% vs. 2.8%, *p* < 0.05), whereas regulatory M2 macrophages (CD206^+^) were decreased in patients with single vessel and multivessel disease (0.34% and 0.34% vs. 1.4%, *p* < 0.05). An inverse relationship between paired CD16^+^ monocytes and M2 macrophages marked CAD severity. CAD was also found to be more tightly associated with CD16^+^ cells than age or traditional cardiovascular risk factors on multiple regression analysis of these patients.

**Conclusions:** CAD extent is correlated directly with CD16^+^ monocytes and inversely with M2 (CD206^+^) macrophages, suggesting circulating monocytes may influence downstream polarization of lesional macrophages. These measures of monocyte and macrophage subtypes hold potential as biomarkers in CAD.

## INTRODUCTION

Atherosclerosis has a significant inflammatory component characterized by an imbalance between pro-inflammatory and regulatory influences [1]. Monocytes and their descendent cells, macrophages, play significant roles in the initiation and progression of the disease [2]. The cells’ involvement in this complex pathophysiology is best understood in the context of cellular subsets, as various classes of monocytes and macrophages serve distinct, characteristic functions [3,4]. Monocytes can be classified as classical, intermediate, or non-classical according to CD14 and CD16 expression. Classical monocytes (CD14^hi^CD16^−^) are involved with phagocytosis and cytokine production. On the other hand, non-classical monocytes (CD14^lo^CD16^hi^) can be pro-inflammatory or potentially pro-angiogenic depending on the context [5,6], and the intermediate monocyte (CD14^hi^CD16^lo^) phenotype partly overlaps with that of non-classical cells [6]. These CD16^+^ cells (i.e. the overlap of intermediate and non-classical monocytes) can be elevated in CAD patients [7,8], and the presence of these cells may associate with plaque vulnerability and rupture [9]. Moreover, intermediate monocytes have been shown to correlate with traditional cardiovascular risk factors and may be predictive of cardiovascular events [10].

Macrophages, derived from all these monocyte subtypes, become the foam cells that comprise the core of the atheroma. They are induced by the local microenvironment to various heterogeneous polarization states along a multidimensional continuum [11,12], but represented at its extremes by so-called “M1” and “M2” macrophages, which can be identified by established cell surface markers and functional phenotype. Classically activated, or M1, macrophages are characterized by expression of pro-inflammatory mediators such as tumor necrosis factor alpha (TNFα) and proteases, which may contribute to plaque destabilization. M2 macrophages, conversely, are anti-inflammatory cells involved in immunoregulation and tissue repair [13]. M1 macrophages are thought to be more prevalent in rupture-prone, unstable regions of atherosclerotic plaques, whereas M2 cells are more prominent in stable plaque regions [14].

Because specific monocyte and macrophage subsets may associate with atherosclerotic burden, characterization of monocyte surface markers and macrophage polarity in CAD may help identify patients with advanced disease who could benefit from more intensive cardiovascular risk reduction. An important step in understanding this relationship is to measure paired monocyte and macrophage phenotypes from patients with differing CAD burden. Comparing matched monocyte and cultured macrophage phenotypes from the same patient would be a novel approach to approximating the fate of circulating monocytes in clinically significant atheroma. Further, if circulating monocyte phenotypes associate with their descendent macrophage polarization states as well as disease burden, these cells could serve as more focused anti-inflammatory therapeutic targets or surrogates than many of those currently under investigation [15]. Therefore, our aim was to characterize the relationship between CAD burden and the matched phenotypes of monocytes and macrophages derived from peripheral blood of patients whose CAD is defined by coronary angiography.

## MATERIALS AND METHODS

### Subjects and study design

Patients scheduled for cardiac catheterization at the University of Chicago were screened for participation in this study. Patients were included if they underwent left heart catheterization and excluded if they had prior heart transplantation. Upon completion of the angiogram, participants were classified according to coronary artery disease (CAD) severity as having no significant CAD, single vessel disease (SVD), or multivessel disease (MVD). The *no CAD* group had no lesions beyond mild or minimal luminal irregularities, whereas the SVD and MVD were classified according to the number of major coronary arteries with either a stent or an atherosclerotic lesion greater than 75% of vessel diameter [16]. The study was approved by the Institutional Review Board of the University of Chicago and conforms to the ethical guidelines of the 1975 Declaration of Helsinki. All participants provided written informed consent.

### Monocyte isolation

2.5 mL of fresh peripheral blood, collected into an anticoagulating citrate tube from the arterial sheath of each patient upon obtaining access, was mixed with an equal volume of phosphate-buffered saline, gently layered over 2.5 mL of Lymphoprep (Axis-Shield, Oslo, Norway), and centrifuged for 20 minutes at 800 × g without brakes. The peripheral blood mononuclear cell (PBMC) layer was collected and washed twice in 3 mL phosphate-buffered saline, centrifuged for 5 minutes at 1500 × g to recover cells after each washing. PBMCs for flow cytometry analysis were harvested after these steps and fixed in 2% paraformaldehyde for 15 minutes, and stored at 4°C. Remaining PBMCs were resuspended in 4 mL RPMI 1640 (Thermo Fisher Scientific, Indianapolis, IN), plated on a 60 mm tissue culture dish for two hours, and allowed to adhere. Media was then aspirated and the dish washed with phosphate-buffered saline to remove non-adherent cells [17].

### Differentiation of monocytes to macrophages and macrophage detachment

Adherent cells were incubated in 4 mL RPMI media with 10% macrophage colony stimulating factor-containing CMG-conditioned media and 1% penicillin/streptomycin [18,19]. Then, 500 μl RPMI supplemented with 10% CMG media, 10% fetal bovine serum (Life Technologies, Carlsbad, CA) and 1% penicillin/streptomycin were added to the media daily for four days, after which the media was replaced with fresh 4 mL RPMI media containing 10% CMG media, 4% fetal bovine serum and 1% penicillin/streptomycin. Macrophages were detached from the dish after 7 days by 15 minute incubation in 2 mL TrypLE (Thermo Fisher Scientific) followed by trituration with RPMI media containing 10% fetal bovine serum. Cells were fixed in 2% paraformaldehyde solution for 15 minutes and stored at 4°C.

### Flow cytometry

Fixed monocytes and macrophages were stained with conjugated monoclonal antibodies (Alexa Fluor 488 CD14 and Alexa Fluor 647 CD16 for monocytes; PE/Cy7 CD86 and Alexa Fluor 488 CD206 for macrophages) (BioLegend, San Diego, CA). Corresponding conjugated isotype antibodies were used in all experiments as negative controls (Alexa Fluor 488 Mouse IgG, clone MOPC-21; Alexa Fluor 647 Mouse IgG, clone MOPC-21; PE/Cy7 Mouse IgG, clone MOPC-11) (BioLegend). A 1:200 dilution was used for both monoclonal and isotype antibodies. Cells were analyzed by flow cytometry (LSR II; BD Biosciences) using FACSDiva software (BD Biosciences). Flow cytometry data were analyzed using Flowing software (University of Turku, Finland).

Monocytes and macrophages were gated in a forward scatter/side scatter plot as described previously [20]. Monocytes were divided into subsets according to CD14 and CD16 expression, with classical monocytes defined as CD14^hi^/CD16^−^, intermediate as CD14^hi^/CD16^lo^, and non-classical as CD14^lo^/CD16^hi^ [21]. Each subset was quantified as a percentage of the total monocytes. Macrophages were classified as M1 or M2 according to CD86 and CD206 positivity, respectively [22]. Each polarization state was quantified as a percentage of the total macrophages.

### Statistical analysis

One-way ANOVA and X^2^ testing was used to compare baseline characteristics. Unpaired, two-tailed Student’s t-tests were used to compare FACS data between groups. Data are presented as mean ± SD. Calculations were performed using GraphPad Prism software (San Diego, CA).

Univariate linear regression was performed using 8 independent variables: Age, WBC count, mean arterial pressure (MAP) just prior to angiogram, body mass index (BMI), serum creatinine, CD16+ monocyte count, M2 macrophage count, and number (ranging from 0 to 5) of cardiovascular risk factors [hypertension, hyperlipidemia, diabetes, smoking, African-American race]. The dependent variable was a validated CAD severity score [23], a simplified alternative to the SYNTAX score [24,25]. The variables significantly associated (*p*<0.05) with CAD severity on univariate analysis were included in multiple linear regression analysis. Analysis was performed with R.

## RESULTS

Samples were obtained from 30 patients, of whom 7 had no significant CAD, 7 had SVD, and 16 had MVD. Baseline characteristics of study participants are shown in **Table 1**. Most measures were similar across groups, but differences were observed in age and statin use which were both greater, as expected, in patients with more severe disease.

**Table 1:** Baseline characteristics of study participants. Data are presented as mean ± SD or count (%). AA, African American; ACE, angiotensin converting enzyme; ARB, angiotensin receptor blocker; BMI, Body Mass Index; CCB, calcium channel blocker; DBP, diastolic blood pressure; EF, ejection fraction; HDL, high-density lipoprotein; LDL, low-density lipoprotein; OT, Other (Pacific Islander, more than one race); SBP, systolic blood pressure; WH, White. *p < 0.05, highlighted in red.

|  | No CAD (n=7) | SVD (n=7) | MVD (n=16) |
| --- | --- | --- | --- |
| **Age (years)\*** | **52.0 ± 10.8** | **65.3 ± 10.0** | **71.0 ± 10.0** |
| **Male sex** | 4 (57%) | 2 (29%) | 11 (69%) |
| **Race** | AA 3 (43%)<br>WH 3 (43%)<br>OT 1 (14%) | AA 5 (71%)<br>WH 2 (29%)<br>OT 0 (0%) | AA 7 (44%)<br>WH 9 (56%)<br>OT 0 (0%) |
| **BMI (kg/m2)** | 30.3 ± 10.8 | 25.9 ± 7.4 | 29.0 ± 4.7 |
| **SBP (mm Hg)** | 127.9 ± 16.7 | 142.7 ± 24.2 | 138.9 ± 25.1 |
| **DBP (mm Hg)** | 78.3 ± 17.5 | 80.7 ± 12.7 | 76.0 ± 17.4 |
| **EF (%)** | 42.3 ± 15.4 | 37.9 ± 21.6 | 42.5 ± 15.0 |
| **Total cholesterol (mg/dl)** | 149.3 ± 23.6 | 179.5 ± 34.1 | 151.9 ± 45.5 |
| **HDL cholesterol (mg/dl)** | 44.0 ± 14.7 | 52.3 ± 17.0 | 42.4 ± 7.4 |
| **Triglycerides (mg/dl)** | 115.0 ± 38.3 | 126.8 ± 24.4 | 131.8 ± 51.8 |
| **LDL cholesterol (mg/dl)** | 82.5 ± 19.8 | 102.3 ± 44.3 | 83.0 ± 38.1 |
| **Leukocytes (1/μl)** | 8.2 ± 3.0 | 8.5 ± 2.4 | 6.4 ± 1.8 |
| **Total monocytes (1/μl)** | 0.54 ± 0.17 | 0.62 ± 0.36 | 0.65 ± 0.30 |
| **% monocytes** | 5.8 ± 2.5 | 7.4 ± 3.0 | 10.3 ± 3.9 |
| **Creatinine (mg/dl)** | 1.0 ± 0.4 | 1.3 ± 0.9 | 1.5 ± 1.4 |
| **Smoking history** | 3 (43%) | 3 (43%) | 9 (56%) |
| **Diabetes Mellitus** | 2 (29%) | 3 (43%) | 4 (25%) |
| **Hypertension** | 5 (71%) | 6 (86%) | 14 (88%) |
| **Hyperlipidemia** | 3 (43%) | 6 (86%) | 11 (69%) |
| **Beta-blockers** | 4 (57%) | 4 (57%) | 15 (94%) |
| **ACE inhibitors** | 3 (43%) | 0 (0%) | 7 (44%) |
| **ARBs** | 0 (0%) | 3 (43%) | 2 (13%) |
| **CCBs** | 0 (0%) | 3 (43%) | 3 (19%) |
| **Antiplatelet drugs** | 7 (100%) | 7 (100%) | 16 (100%) |
| **Statins\*** | **4 (57%)** | **6 (86%)** | **16 (100%)** |

To assess circulating monocyte phenotypes, each patient’s PBMCs were isolated from fresh blood and stained for monocyte markers, then measured by flow cytometry with gating on the monocyte population and quantification of CD14 and CD16 expression. Classical monocytes were the majority of circulating monocytes for all patients and did not show a consistent pattern as a function of CAD severity (**Fig 1A**). Although they remained a minority population, non-classical monocytes were increased in patients with SVD (5.2±1.1 vs. 1.5±0.2%; *p* < 0.01) and MVD (6.8±1.4 vs. 1.5±0.2%; *p* < 0.05) compared to those without significant CAD (**Fig 1B**). Intermediate monocytes also showed this trend (compared to *no CAD*, p = 0.08 for SVD and 0.05 for MVD, **Fig 1C**). Total CD16^+^ (combined non-classical and intermediate) monocytes were similarly elevated in SVD (8.6±1.8 vs. 2.8±0.6%; *p* < 0.05) and MVD (10.5±1.7 vs. 2.8±0.6%; *p* < 0.01), both in absolute percentages (**Fig 1D**) and when expressed relative to classical monocytes (**Fig 1E**).

**Figure 1:**
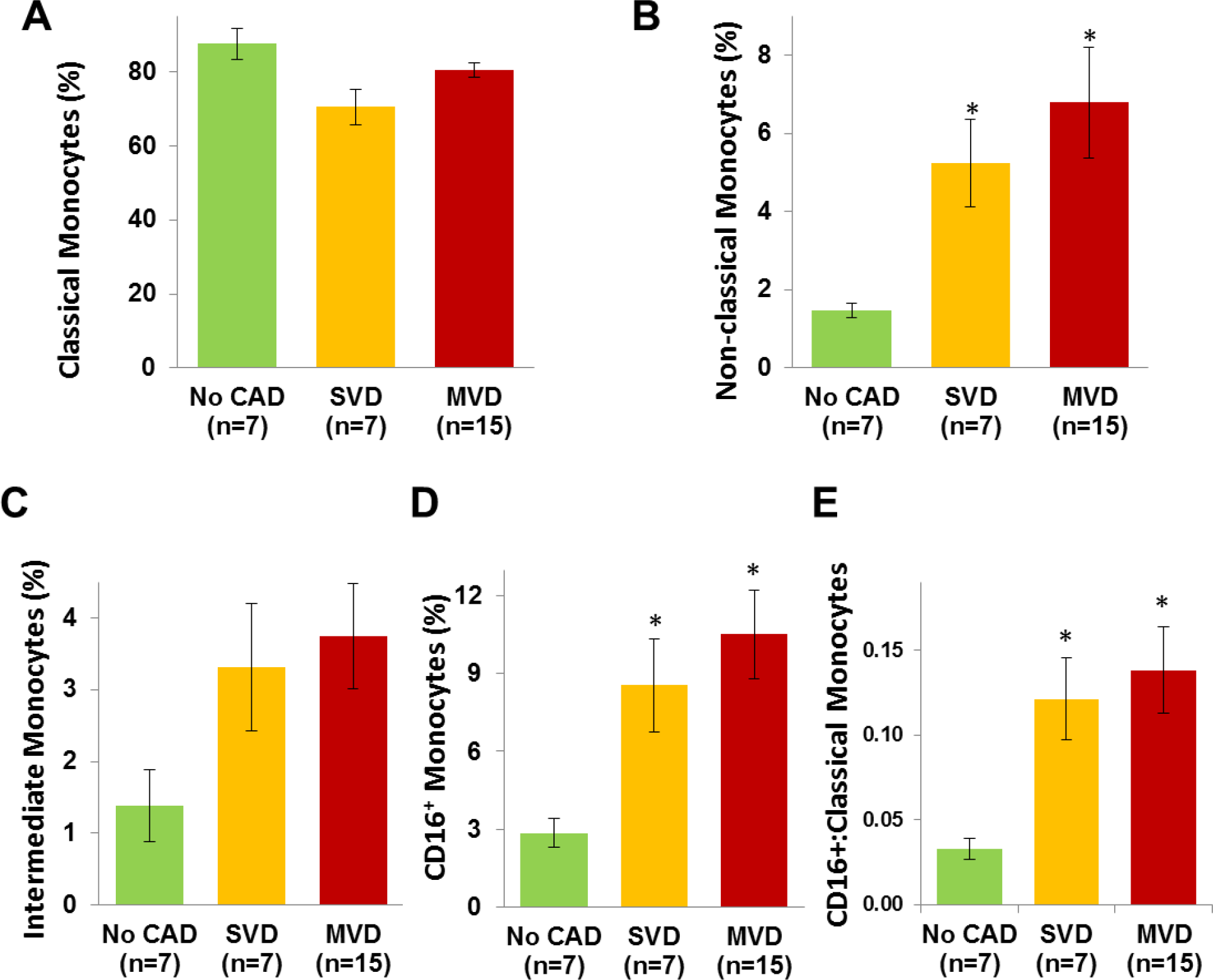
Monocytes subtypes as a function of CAD burden. (**A**) Classical monocytes are the majority of circulating monocytes and show no consistent pattern as a function of CAD severity. (**B**) Although still a minority population, non-classical monocytes were relatively enriched in patients with CAD. (**C**) Intermediate monocytes also showed this trend (compared to *no CAD*, p = 0.08 for SVD and 0.05 for MVD). (**D**) Total CD16^+^ monocytes (intermediate and non-classical combined) are shown, as is the ratio of CD16^+^ to classical monocytes (**E**), all as a function of CAD burden. Data are presented as mean ± SEM. * p < 0.05 in all panels.

To determine whether differences in circulating monocyte phenotype translate into differences in macrophage polarization, each patient’s monocytes, cultured in conditions that drive macrophage differentiation [18,19], were differentiated over seven days into adherent peripheral blood-derived macrophages. These cells were assessed for macrophage polarization markers CD86 and CD206 by flow cytometry. No significant differences in M1 macrophages (CD86^+^/CD206^−^) were detected as a function of CAD burden (**Fig 2A**), but M2 macrophages (CD86^−^/CD206^+^) were decreased in patients with SVD (0.3±0.1 vs. 1.4±0.4%; *p* < 0.05) and MVD (0.3±0.1 vs. 1.4±0.4%; *p* < 0.01) compared to those without significant CAD (**Fig 2B**). The ratio of M1:M2 macrophages was elevated in SVD (360±92 vs. 125±45; *p*<0.05) and MVD (434±73 vs. 125±45; p<0.05) (**Fig 2C**). Regarding a potential relationship between distribution of monocyte subsets matched to their descendent macrophage polarization, CAD severity roughly fell on an inverse relationship between CD16^+^ monocytes and differentiation into M2 macrophages (**Fig 2D**).

**Figure 2:**
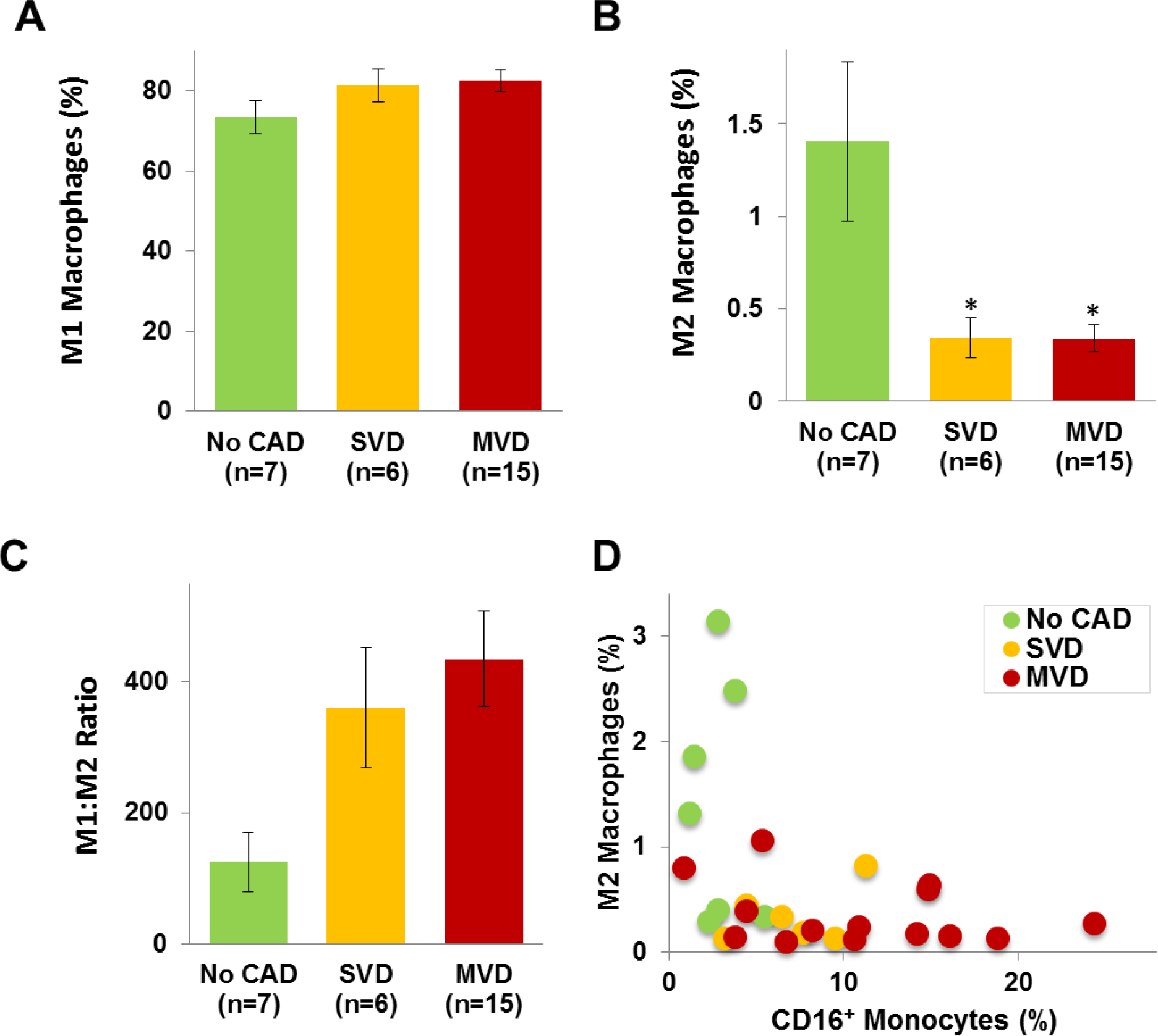
Profiles of cultured macrophage from patients with CAD. (**A**) Of the macrophages cultured from all patients with or without CAD, most had the M1 surface marked (CD86), without significant differences between clinical presentations. (**B**) The minority population of M2 macrophages (CD206^+^) was suppressed in patients with CAD. (**C**) The M1:M2 macrophage ratio rises according to CAD severity. Data are presented as mean ± SEM. *p < 0.05. (**D**) M2 macrophages plotted against CD16^+^ monocytes show an inverse relationship that tracks with CAD burden.

Univariate linear regression was performed using several documented clinical variables, in addition to CD16^+^ and M2 counts, to determine association with CAD severity based on an established score that serves as a simplified alternative to the SYNTAX score [23]. CD16^+^ monocyte count, age, and total number of cardiovascular risk factors [hypertension, hyperlipidemia, diabetes, smoking, African-American race] were significantly and directly associated with CAD severity, whereas M2 macrophage count was inversely associated with CAD severity (**Table 2**). When these four variables were analyzed through multiple regression, CD16^+^ monocytes remained significantly associated with CAD severity (**Table 2**).

**Table 2:** Clinical metrics and monocyte/macrophage markers associated with CAD severity. Univariate linear regression was performed using 8 independent variables in association with each patient’s CAD severity score. Standardized correlation coefficients (β) are shown. The variables significantly associated (p < 0.05) with CAD severity were CD16^+^ monocytes, age, cumulative risk factors, and M2 macrophages (inversely). These were included in multiple linear regression analysis, in which CD16^+^ count was the most significant association.

|  | Univariate |  | Multivariate |  |
| --- | --- | --- | --- | --- |
| | $\beta$ | p | $\beta$ | p |
| **CD16<sup>+</sup> Monocytes** | **0.64** | **0.0003** | **0.43** | **0.01** |
| **Age** | **0.59** | **0.001** | 0.26 | 0.14 |
| **Risk Factors** | **0.5** | **0.01** | 0.23 | 0.18 |
| **M2 Macrophages** | **-0.43** | **0.03** | -0.11 | 0.48 |
| **WBC** | -0.34 | 0.08 |  |  |
| **Creatinine** | 0.32 | 0.11 |  |  |
| **MAP** | -0.16 | 0.42 |  |  |
| **BMI** | -0.08 | 0.68 |  |  |

## DISCUSSION

A major limitation in understanding the pathogenesis of CAD is the inability to study living cells from the coronary plaques of living patients. Surrogate biomarkers abound, but a cell biomarker based on monocytes and macrophages, main participants in plaque initiation and growth, may help to clarify CAD progression and risk further. In this study, we characterized monocyte subsets matched with their *ex vivo* macrophage profiles from peripheral blood samples of patients with varying CAD severity. Non-classical and total CD16^+^ monocytes were elevated in CAD patients, and these patients’ monocytes, once differentiated into macrophages, demonstrated impaired expression of markers associated with a regulatory, anti-inflammatory state. CD16^+^ monocytes and M2 macrophages thus showed inverse correlation with each other, suggesting that circulating monocytes influence polarization of their descendant macrophages in target tissues, such as the coronary atheroma. This is a new approach to systematically culture peripheral blood-derived macrophages for the purpose of comparing monocyte and monocyte-derived macrophage phenotypes from the same patient and correlating to his or her CAD burden.

Our finding that non-classical monocytes are increased in stable CAD is consistent with previous studies. While all three monocyte subsets play a role in the initiation and progression of the atheroma, non-classical monocytes are distinguished by their capacity for recruitment to inflamed tissues [6]. In addition to CD16, these cells express CX3CR1, the receptor for fractalkine, an adhesion molecule expressed on activated endothelium [26]. Thus, an increase in the proportion of non-classical monocytes may be associated with progression of stable CAD through recruitment of these cells to the intima [27]. Additionally, the decrease in M2 macrophages and the imbalance between M1 and M2 phenotypes we observe is consistent with findings in mouse models of atherosclerosis. These murine studies suggest that M2 macrophages dominate during the initiation of the atherosclerotic lesion, whereas a shift to the M1 phenotype is observed with disease progression [28]. The shift away from regulatory M2 macrophages that we measure could reflect a transition from early to established disease, consistent with these patients’ clinical status. Further, the concept of a skewed ratio of pro-inflammatory and regulatory macrophages predisposing to atherosclerosis has been backed by murine and human studies [13,29,30].

Several limitations have bearing on the interpretation of our data. Exclusion criteria were kept to a minimum to allow for a range of patients with multiple comorbidities, which likely contributed to the variability in data, yet this better reflects real clinical practice.
The small sample size also limits our statistical power. Finally, differences in other clinical factors between the groups may have an impact on the expression of these biomarkers, although many, including age and traditional risk factors, were taken into account for the multivariable analysis.

In sum, non-classical and CD16^+^ monocytes are increased and monocyte-derived M2 regulatory macrophages are decreased in patients with CAD compared to those without. That the differences in macrophage polarization persist after a week of *ex vivo* culture demonstrates a durability in the impact of monocyte phenotype on eventual macrophage polarization, a hypothesis supported by studies of monocyte epigenetic memory [31]; this suggests that both circulating monocytes and tissue macrophages may be useful therapeutic targets for preventing initiation and progression of disease. Indeed, translational studies examining the pleotropic effects of cardiovascular medications on monocytes and macrophages have demonstrated changes in these cells’ phenotype and function in response to treatment [32–38]. Further, distributions of pro-inflammatory monocytes and macrophages hold potential as markers of stable CAD burden. In our data, CD16^+^ monocytes in particular were more closely associated with quantified CAD burden than traditional cardiovascular risk factors. As alterations in monocyte and macrophage profiles have been identified in human studies of carotid and femoral plaque rupture [39–41], our methods may allow for future replication of these findings in surgically excised coronary plaques. Finally, future studies in patients with unstable CAD could help determine whether pro-inflammatory monocytes and macrophages can better predict acute coronary syndromes.

